# Modulatory Effect of Levodopa on the Basal Ganglia-Cerebellum Connectivity in Parkinson’s Disease

**DOI:** 10.1101/2023.01.16.524229

**Authors:** Subati Abulikemu, Yen F Tai, Shlomi Haar

## Abstract

Levodopa has remained the mainstay of medical therapy for Parkinson’s disease since its development in the 1980s. However, long-term medication use is associated with declining clinical efficacy and the emergence of motor complications. Unveiling the effects of levodopa on brain functional reorganisation at a relatively early treatment phase is therefore imperative to inform the optimisation of Parkinson’s therapeutics. In this study, we comprehensively investigated levodopa’s modulation on the resting-state functional connectivity in the cortico-basal ganglia-cerebellum system at regional and network levels, with dual cross-sectional and longitudinal designs. The data was extracted from the Parkinson’s Progression Marker Initiative (PPMI) dataset. The cross-sectional patient groups comprised 17 Parkinson’s patients on stable levodopa medication and 15 drug-naïve patients, while the longitudinal set included 14 Parkinson’s patients measured at both drug-naïve and levodopa-medicated conditions. With nodes defined across cortical, basal ganglia, and cerebellar networks, we conducted univariate comparisons of the internodal connectivity strength between the medication conditions using nonparametric permutation. At the network level, we computed multivariate combinations of individual connections within and between the networks, followed by an assessment of their discriminative capabilities on patients’ medication classes using supervised machine learning. The univariate seed-based approach showed no statistically significant effect of levodopa in either dataset. However, the network connectivity pattern between basal ganglia and the cerebellum displayed a robust classification power in the longitudinal dataset and a similar trend was observed in the cross-sectional. The role of the cerebellum is often overlooked in previous functional integration investigations of Parkinson’s disease and levodopa effects. Considering the recent evidence suggesting the bidirectional communications between the cerebellum and basal ganglia networks, our study provides further insight into the importance of inter-network functional connectivity in Parkinson’s, as well as in the functional and plastic processes following levodopa medication.

## Introduction

Parkinson’s disease (PD) is the second most prevalent neurodegenerative disorder affecting 1- 2% of the population over 50 years of age (Shastry, 2001). Clinically, PD is a movement disorder characterised by bradykinesia, resting tremor, rigidity, and postural instability (Haddad et al., 2018; Shastry, 2001). The cardinal pathology of PD is the depletion of nigrostriatal dopaminergic (DA) neurons with consequent dysfunction of cortico-striatal-thalamic-cortical circuits(Braak et al., 2004; Hacker et al., 2012) (CSTC). Since its development in the late 1960s, levodopa (L-3,4-dihydroxyphenylalanine), as a direct precursor to DA, has remained the most efficacious symptomatic therapy for PD (Poewe and Espay, 2020). With prolonged treatment, however, the clinical response to levodopa progressively declines (i.e., wearing-off phenomenon) and leads to the occurrence of motor complications, including fluctuations, dyskinesia, and dystonia (Mueller et al., 2019; Thanvi and Lo, 2004). Hence, it is crucial to uncover levodopa-induced neural effects and functional brain reorganisation to provide new insights into optimising PD therapeutics.

Resting-state functional connectivity (rs-FC) has been increasingly examined in PD brains and their modulations following levodopa medication, in accordance with the nexopathy framework conceptualising neurodegenerative diseases as disconnection syndromes (Warren et al., 2013). Aberrant functional integration between cortical sensorimotor areas and striatum was consistently detected and deemed a fundamental pathological remapping in PD (Helmich et al., 2010; Luo et al., 2014; Tahmasian et al., 2015). Levodopa has been shown to modulate the functional coherence in the basal ganglia (BG)-thalamic-motor cortical system, including conflicting down- and up-regulated connectivity changes, potentially due to heterogeneity in methodology, clinical states of patients, and seed selections (Tahmasian et al., 2015). Past reports were often based on the acute levodopa responses, and longitudinal rs-FC changes following stable levodopa medication in drug-naïve patients are yet to be extensively investigated.

Apart from the conventional CSTC circuit dysfunction in PD, the role of the cerebellum has been emphasised. The cerebellum is likely to exert both crucial pathological and compensatory effects in PD (Wu and Hallett, 2013). Relative to healthy controls, PD patients displayed weakened striatum-cerebellum connectivity, significantly greater activations of the bilateral cerebellum, and strengthened functional coupling between the cerebellum and the cortical motor network (Bagarinao et al., 2022; Jahanshahi et al., 2010; Wu et al., 2011). The diminished FC might reflect the aberrant BG signalling over the cerebellum in PD, whereas the cerebellar-thalamic-cortical circuit was thought to be increasingly strengthened as the pathology progresses to preserve motor functions (Bostan and Strick, 2010; Wu and Hallett, 2013). Acute levodopa administration was shown to substantially elevate the cerebellum connectivity to subcortical regions of the motor system, including the thalamus, putamen, globus pallidus (GP), and brainstem (Mueller et al., 2019). A recent consensus paper emphasised the role of the cerebellum within the integrated cortico-BG (-thalamo)-cerebellum system where functional and plasticity processes of the networks were interactive (Caligiore et al., 2016). A system-level mechanism of cerebellum, cortex, and BG in Parkinson’s and its plastic changes following levodopa medication is yet to be further examined.

In this project, we adopted both cross-cohort and longitudinal designs to investigate the effect of levodopa on motor-functional reorganisation in PD under stable levodopa treatment. The former entailed two independent groups of de novo and levodopa-medicated patients with matching disease durations, whereas the latter included a single PD cohort who transitioned from drug-naïve to levodopa-medicated states. We investigated regional and network connectivity in the motor cortex, BG, and cerebellum employing respective seed- and network-level analyses. Our approach enabled the assessment of levodopa modulation at varying neural levels and the comparisons of which under different research designs.

## Results

### 1.1 Sample selection

The longitudinal cohort consisted of 14 patients who had resting-state scans at both de novo and levodopa-medicated states (Table 2). Given that 10 patients had multiple scans while on levodopa medication, we determined a combination of levodopa scans that produced the least variance in treatment duration (Figure 1A). To derive cross-sectional de novo and levodopa groups with comparable disease spans, we first inspected the distribution of disease durations at all 184 resting-state (rs)-functional Magnetic Resonance Imaging (fMRI) visits (73 de novo and 111 levodopa scans; Figure 1B). A cut-off of 15-40 months was then applied, and the disease duration more proximal to the group mean was selected if the subject attended multiple scanning sessions. For chosen levodopa patients, medication duration was further filtered with a threshold of 12-24 months. The procedure resulted in 15 de novo and 17 levodopa patients (Table 2), with the de novo group manifesting a relatively more diffused and multimodal distribution of disease duration (Figure 1C). Patients were ON-medication at all levodopa scans, with *median* time elapsed since the last intake of 3.34 hours [Interquartile Range (*IQR*) = 2.91 hours] and 2.99 hours (*IQR* = 2.01 hours) for longitudinal and cross-sectional datasets, respectively.

**Figure 1.**
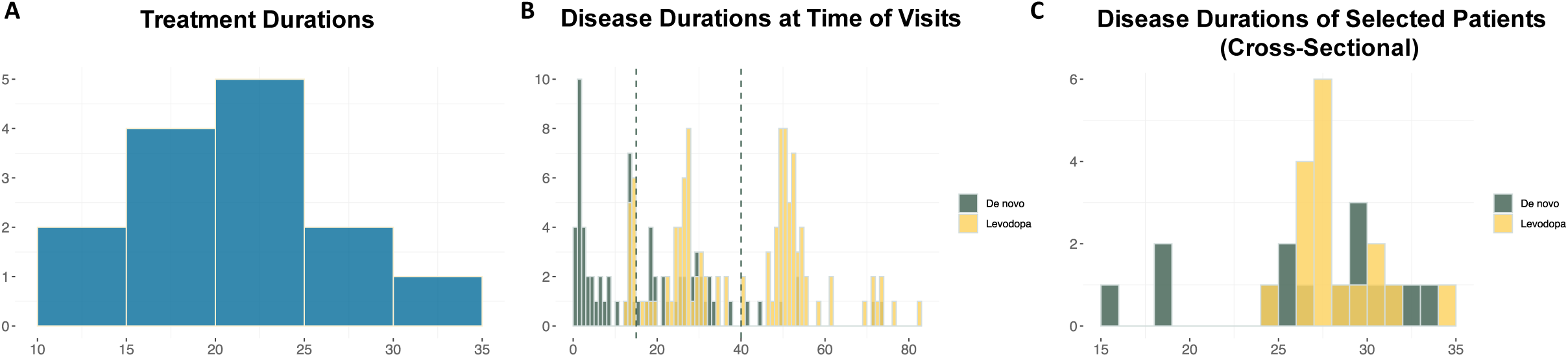
Sample selection based on treatment and disease durations. (**A)** Longitudinal cohort: treatment durations at selected levodopa scans (*median* = 22 months; *IQR* = 6.75 months; *maximum* = 35 months; *minimum* = 13 months). (**B)** The distribution of disease durations at all rs-fMRI visits (de novo: *median* = 14 months; *IQR* = 25 months; *maximum* = 74 months; *minimum* = 1 months; levodopa: *median* = 47 months; *IQR* = 25 months; *maximum* = 83 months; *minimum* = 13 months). The dashed lines represent the threshold applied for the selection of the cross-sectional cohort. **(C)** Cross-sectional cohort: the spreading of disease durations after the filtering and the removal of missing scans (de novo: *median* = 29 months; *IQR* = 5 months; *maximum* = 34 months; *minimum* = 15 months; levodopa: *median* = 28 months; *IQR* = 3 months; *maximum* = 35 months; *minimum* = 25 months).

Table 2 provides the demographic and clinical descriptors of the selected patients. In the longitudinal dataset (Table 2A), the median interval between two visits was 34.50 months (*IQR* = 11 months). At time point two, ON-levodopa patients (within 6 hours after the last levodopa intake) experienced a significant attenuation of motor symptoms measured by part three of the Movement Disorder Society-Unified Parkinson’s Disease Rating Scale (UPDRS-III; Goetz et al., 2008) (*median of paired difference* = 6.50, *IQR* = 8.50, *p* < 0.01). The decreased score was driven by bradykinesia/rigidity (*median* = 2.50, *IQR* = 6.50, *p* = 0.02) and tremor (*median* = 3, *IQR* = 5, *p* = 0.02) measures. At the same time, when tested as the medication wears off (over 6 hours after the last levodopa intake) there was no significant change in motor symptoms from the de novo measures taken about three years earlier. In the cross-cohort set (Table 2B), De novo and medicated patients were matched for demographic variables. The medicated patients displayed reduced UPDRS-III scores (*difference between medians* = 10; *p* < 0.01) and ameliorated severity of bradykinesia/rigidity (*difference between medians* = 4; *p* = 0.04) while on medication (within 6 hours after the last levodopa intake). UPDRS-III scores after the medication wears off (over 6 hours after the last levodopa intake) were not recorded for this group.

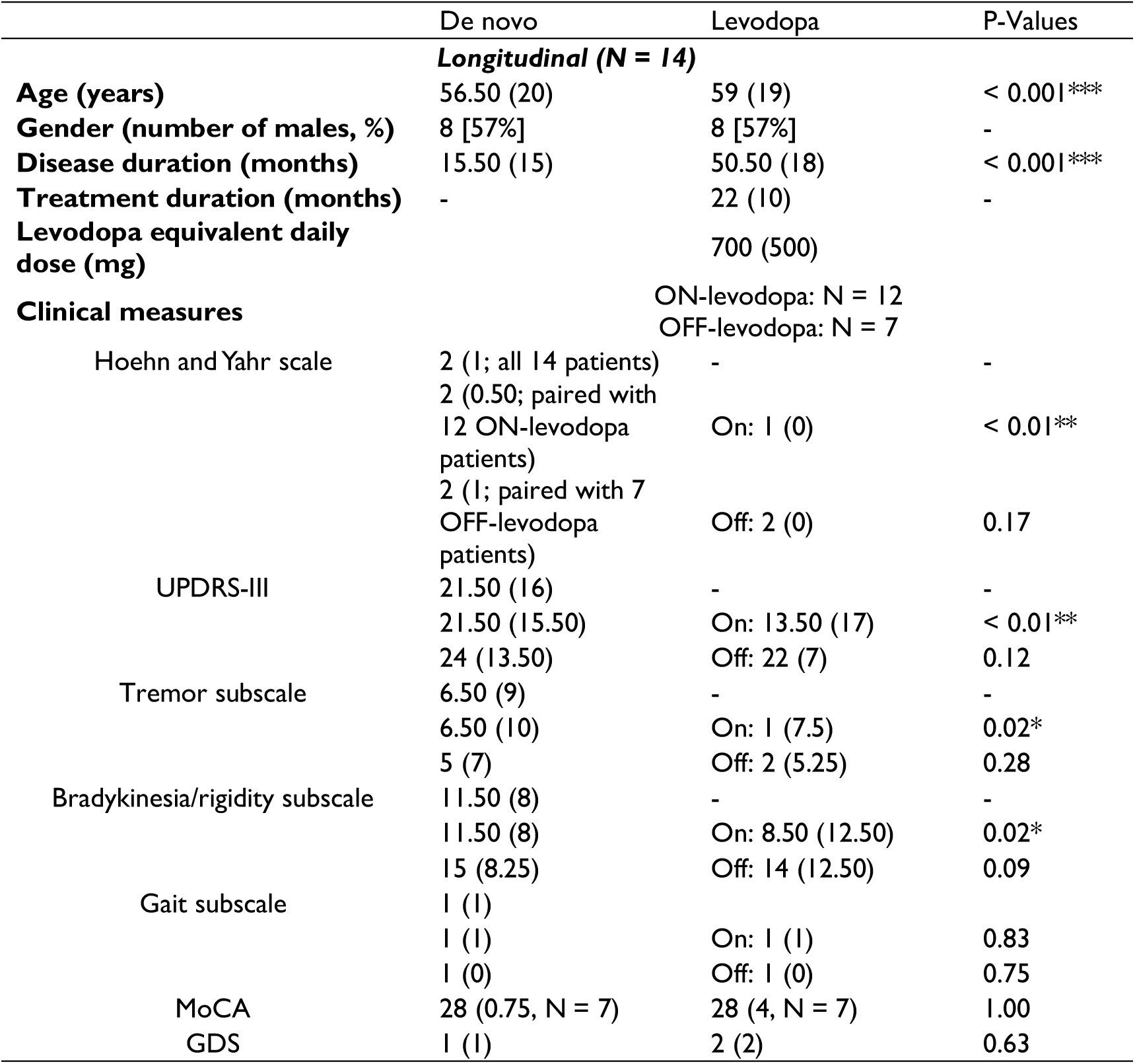
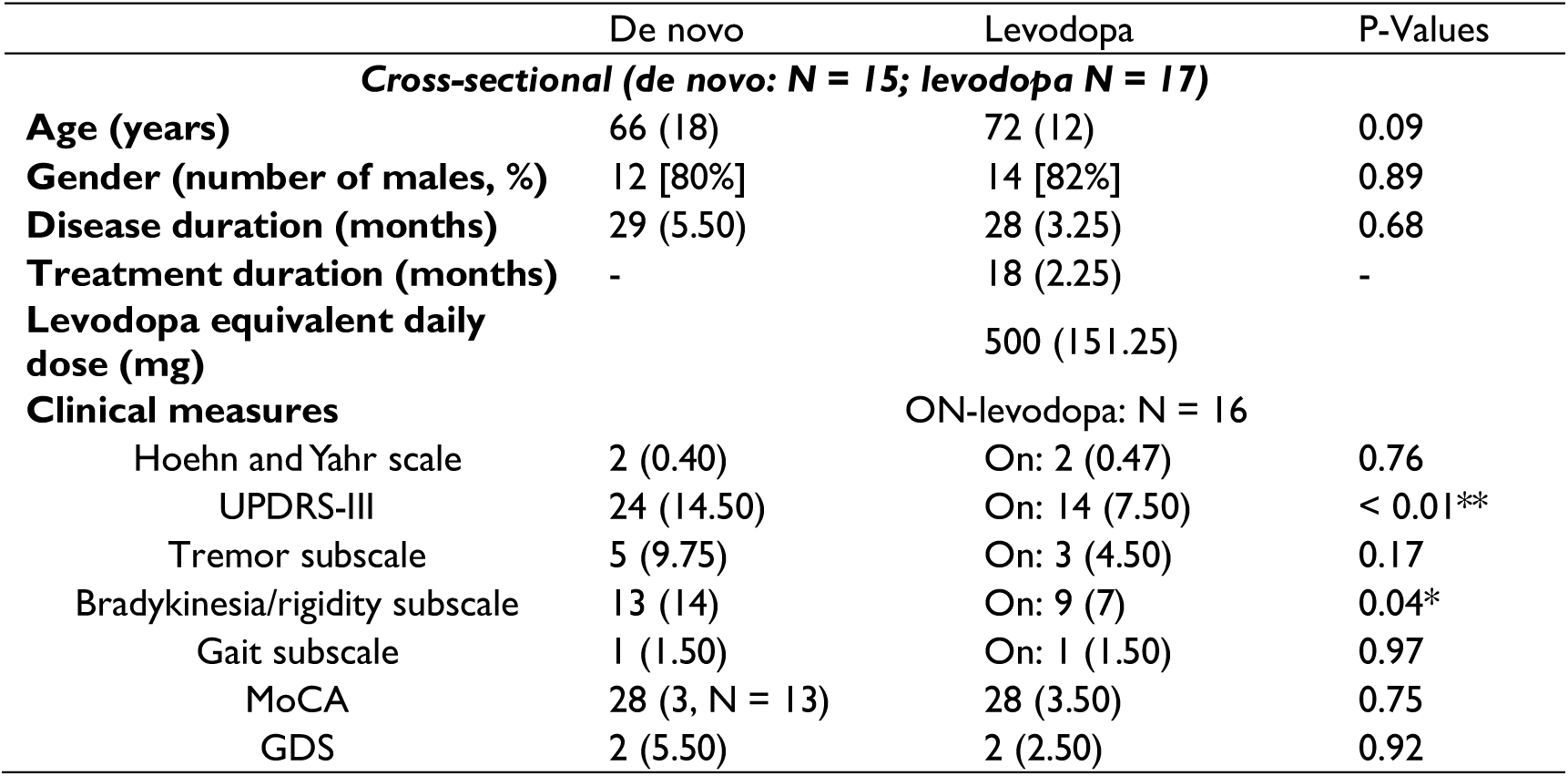
**A. Demographics and clinical measures of patients in the longitudinal dataset.** **B. Demographics and clinical measures of patients in the cross-sectional dataset.** Demographics and clinical measures of study participants. **A**: Longitudinal cohort. **B**: Cross-sectional cohort. Values are medians (interquartile range). For gender, values are numbers of males [% of males]. For patients who initiated levodopa treatment, the OFF levodopa was defined as not having taken medication for at least 6 hours. We performed paired-sample comparisons of the UPDRS measures between the de novo state with ON (N = 12) and OFF (N = 7) levodopa states for the longitudinal dataset. However, only ON (N = 16) scores were included for the cross-cohort comparison, as only 4 levodopa-medicated patients had OFF scores. Moreover, the ON/OFF levodopa were not distinguished for MoCA and GDS. MoCA scores of 7 patients in the longitudinal dataset and 2 patients in the de novo group of the cross-sectional dataset were missing. Due to the non-Gaussian distributions of the variables, we conducted non-parametric Wilcoxon signed-rank and rank-sum tests for within- and between-group comparisons, respectively.

|  | De novo | Levodopa | P-Values |
| --- | --- | --- | --- |
| **Longitudinal (N = 14)** |  |  |  |
| **Age (years)** | 56.50 (20) | 59 (19) | < 0.001*** |
| **Gender (number of males, %)** | 8 [57%] | 8 [57%] | - |
| **Disease duration (months)** | 15.50 (15) | 50.50 (18) | < 0.001*** |
| **Treatment duration (months)** | - | 22 (10) | - |
| **Levodopa equivalent daily dose (mg)** |  | 700 (500) |  |
| **Clinical measures** |  | ON-levodopa: N = 12<br>OFF-levodopa: N = 7 |  |
| Hoehn and Yahr scale | 2 (1; all 14 patients) | - | - |
|  | 2 (0.50; paired with 12 ON-levodopa patients) | On: 1 (0) | < 0.01** |
|  | 2 (1; paired with 7 OFF-levodopa patients) | Off: 2 (0) | 0.17 |
| UPDRS-III | 21.50 (16) | - | - |
|  | 21.50 (15.50) | On: 13.50 (17) | < 0.01** |
|  | 24 (13.50) | Off: 22 (7) | 0.12 |
| Tremor subscale | 6.50 (9) | - | - |
|  | 6.50 (10) | On: 1 (7.5) | 0.02* |
|  | 5 (7) | Off: 2 (5.25) | 0.28 |
| Bradykinesia/rigidity subscale | 11.50 (8) | - | - |
|  | 11.50 (8) | On: 8.50 (12.50) | 0.02* |
|  | 15 (8.25) | Off: 14 (12.50) | 0.09 |
| Gait subscale | 1 (1) |  |  |
|  | 1 (1) | On: 1 (1) | 0.83 |
|  | 1 (0) | Off: 1 (0) | 0.75 |
| MoCA | 28 (0.75, N = 7) | 28 (4, N = 7) | 1.00 |
| GDS | 1 (1) | 2 (2) | 0.63 |
| **Cross-sectional (de novo: N = 15; levodopa N = 17)** |  |  |  |
| **Age (years)** | 66 (18) | 72 (12) | 0.09 |
| **Gender (number of males, %)** | 12 [80%] | 14 [82%] | 0.89 |
| **Disease duration (months)** | 29 (5.50) | 28 (3.25) | 0.68 |
| **Treatment duration (months)** | - | 18 (2.25) | - |
| **Levodopa equivalent daily dose (mg)** |  | 500 (151.25) |  |
| **Clinical measures** |  | ON-levodopa: N = 16 |  |
| Hoehn and Yahr scale | 2 (0.40) | On: 2 (0.47) | 0.76 |
| UPDRS-III | 24 (14.50) | On: 14 (7.50) | < 0.01** |
| Tremor subscale | 5 (9.75) | On: 3 (4.50) | 0.17 |
| Bradykinesia/rigidity subscale | 13 (14) | On: 9 (7) | 0.04* |
| Gait subscale | 1 (1.50) | On: 1 (1.50) | 0.97 |
| MoCA | 28 (3, N = 13) | 28 (3.50) | 0.75 |
| GDS | 2 (5.50) | 2 (2.50) | 0.92 |

### 1.2 Functional connectivity strength mapping

The general connectivity patterns (Figure 2) across the subregions of cortical motor areas, BG, and cerebellum were similar between de novo and levodopa states across the two datasets. Strong interconnectedness within the cortical motor network and cerebellum, moderate connections within BG and thalamus, and limited cross-network coherence were commonly manifested.

**Figure 2.**
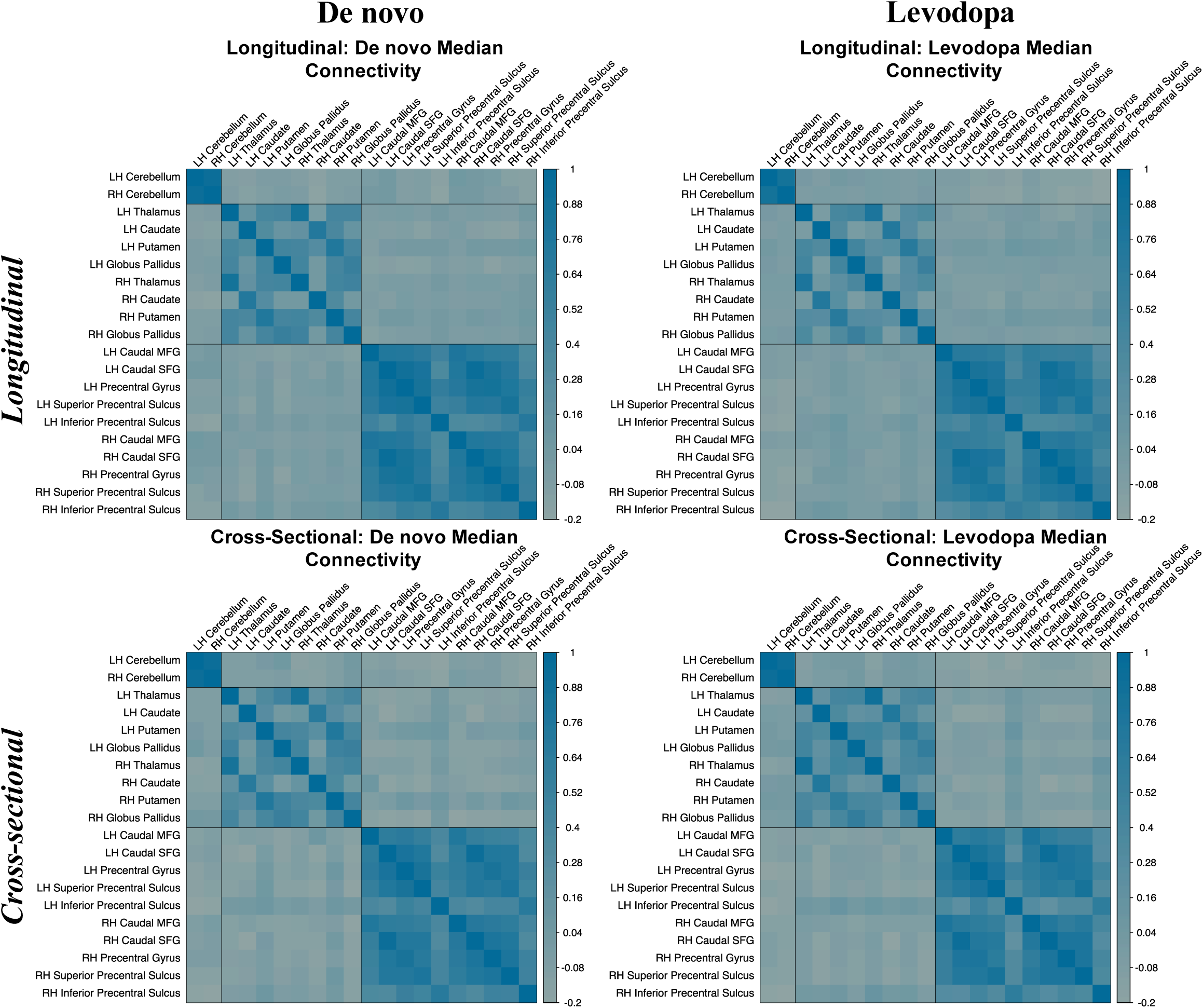
Median interregional functional connectivity strength. The columns show de novo and levodopa medication states and rows include longitudinal and cross-sectional datasets. Within each connectivity matrix, median strength of each interregional rs-FC within each medication condition are represented. Regions within each matrix are organised in the order of cerebellar, basal ganglia (and thalamus), and motor cortical regions. LH = left hemisphere; RH = right hemisphere; MFG = middle frontal gyrus; SFG = superior frontal gyrus.

### 1.3 Within- and between-group comparison of connectivity strength

We first statistically assessed the univariate effect of levodopa on each interregional functional coupling across the three networks. In the longitudinal group, the distribution of p-values in within-subject comparisons of connectivity strength was right-skewed (Figure 3A) and displayed a meaningful trend towards significance. Before correcting for the multiple comparisons, the univariate tests showed higher cerebellum-motor cortex, within-BG, and within-motor cortex functional synchrony at the drug-naïve state and stronger putamen- cerebellum and GP-motor cortex connectivity following the levodopa medication (Figure 3C) under a significance cut-off of 0.05. Nevertheless, no region-of-interest (ROI)-level comparisons survived false discovery rate (FDR) correction. In the cross-sectional dataset, we found no evidence showing differences in interregional connectivity between de novo and levodopa patients. The permutation testing yielded a uniform distribution of p-values (Figure 3A), suggesting that all univariate null hypotheses were true. Hence, the significant tests (before multiple comparisons correction) which showed heightened thalamic-cortical and bilateral thalamic connectivity in the levodopa group and more intense interhemispheric connectivity of areas in the primary motor cortex (Figure 3B) in the de novo group were most likely type I errors.

**Figure 3.**
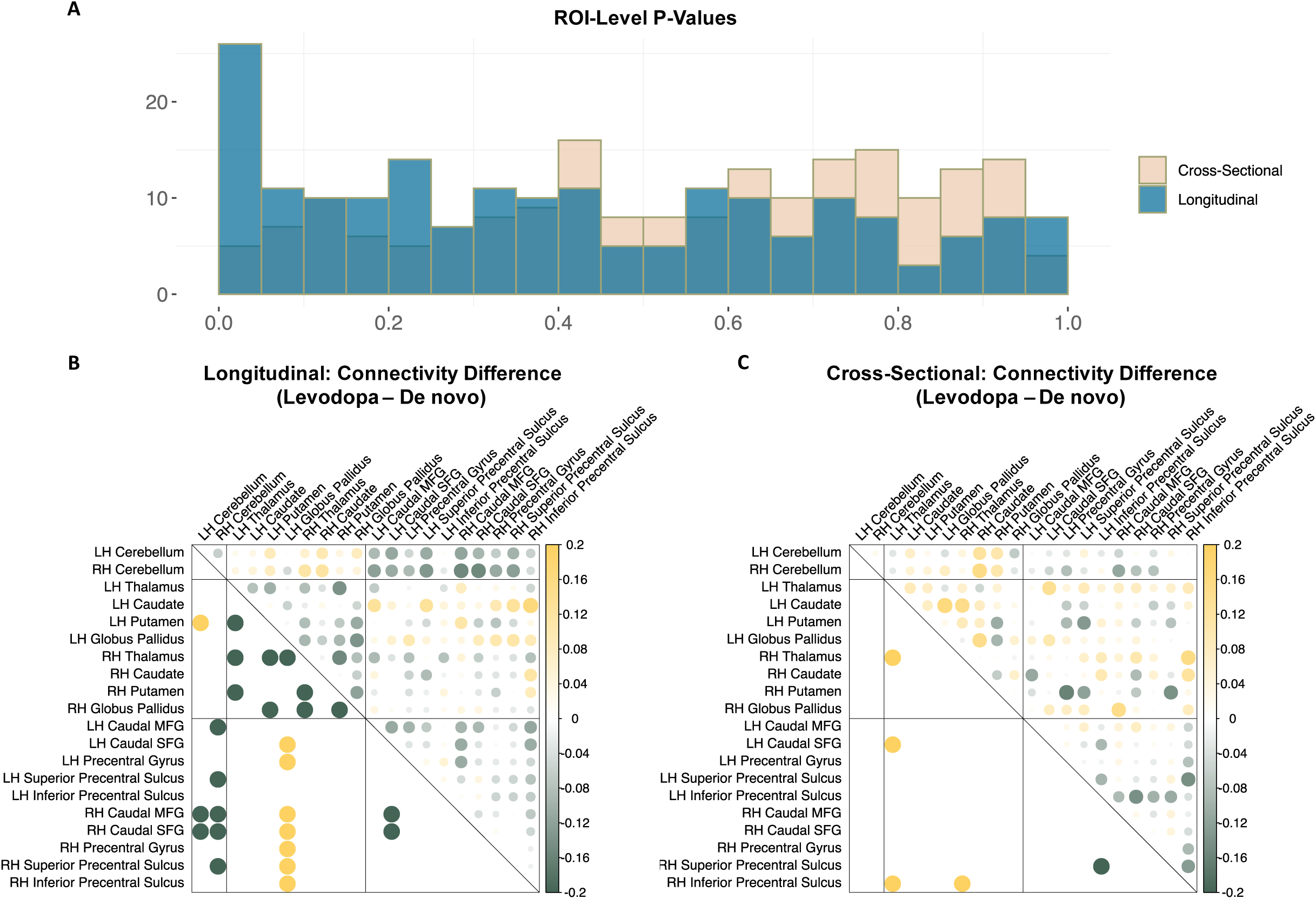
Within- and between-group comparisons in region-of-interest analysis. (A) Frequency histograms of p-values in region-of-interest analyses in cross-sectional (cream) and longitudinal (blue) datasets. **(B) Upper triangle:** Median differences of connectivity magnitude between levodopa and de novo states in longitudinal analysis; size of the circles is proportional to the absolute values. **Lower triangle:** Significant within-subject connectivity contrasts at p < 0.05 prior to false discovery rate correction (yellow circles represent the levodopa connections greater than de novo, and green circles depict more intense de novo connections relative to levodopa). **(C) Upper triangle:** Contrasts of median connectivity strength between levodopa and de novo patients in cross-sectional analysis. **Lower triangle:** Significant between-group connectivity comparisons at p < 0.05 before false discovery rate correction.

### 1.4 Network level differences in functional connectivity

To test the potential effect of levodopa medication on functional connectivity on a network level, i.e., beyond individual ROI-to-ROI connections, we used a support vector machine (SVM) to classify the patterns of connectivity in the network. We considered three subnetworks of the motor CSTC circuitry (cortical motor areas, BG, and cerebellum) and classified the connections within them and between each pair.

The linear SVM implemented with the BG-cerebellum feature set yielded the highest discriminating efficacy in both datasets. In the longitudinal group, the model accurately performed 11 out of 13 classifications of levodopa cases (*sensitivity* = 0.85) and 12 out of 15 predictions of a drug-naïve state (*specificity* = 0.80). The classifier reached an overall *accuracy* of 82.14% and an *area under the receiver operating characteristic curve (AUC)* of 0.80 (Figures 4 & 5). The permutation test showed that the accuracy is significantly higher than chance (*p* = 0.003), and the significance was retained after adjusting for multiple testing (*p* = 0.014). The model fitted with within-BG features had a moderate *accuracy* of 60.71% with a *sensitivity* of 0.59 and a *specificity* of 0.64; however, the accuracy level was not higher than chance (Figure 4; *uncorrected p* = 0.162; *FDR-adjusted p* = 0.350). There was no evidence suggesting the discriminative capacity of the cerebellum-motor cortex (*accuracy* = 0.54; *uncorrected p* = 0.350; *FDR-adjusted p* = 0.350), BG-motor cortical (*accuracy* = 0.57; *uncorrected p* = 0.220; *FDR-adjusted p* = 0.350), or within-motor cortical (*accuracy* = 0.54; *uncorrected p* = 0.315; *FDR-adjusted p* = 0.350) feature sets on patients’ medication class. In the cross-sectional group, the SVM model trained with BG-cerebellum features achieved an *accuracy* of 0.75 and an *AUC* of 0.69 (Figures 4 & 5), where 12 out of 15 patients that were classified into the levodopa group were true positives (*sensitivity* = 0.80) and 12 out of 17 de novo predictions were true negatives (*specificity* = 0.71). Although the observed accuracy value exceeded the 97.5th percentile of the null distribution (*uncorrected p* = 0.012), the statistical significance did not remain after the multiple testing adjustment (*adjusted p* = 0.062). Similar to the longitudinal set, within-BG (Figure 4; *accuracy* = 0.53; *uncorrected p* = 0.317; *FDR-adjusted p* = 0.396), cerebellum-motor cortex (*accuracy* = 0.59; *uncorrected p* = 0.189; *FDR-adjusted p* = 0.339), BG-motor cortical (*accuracy* = 0.47; *uncorrected p* = 0.464; *FDR-adjusted p* = 0. 464), and within-motor cortical (*accuracy* = 0.56; *uncorrected p* = 0.203; *FDR-adjusted p* = 0. 339) feature sets did not manifest significant discriminative power.

**Figure 4.**
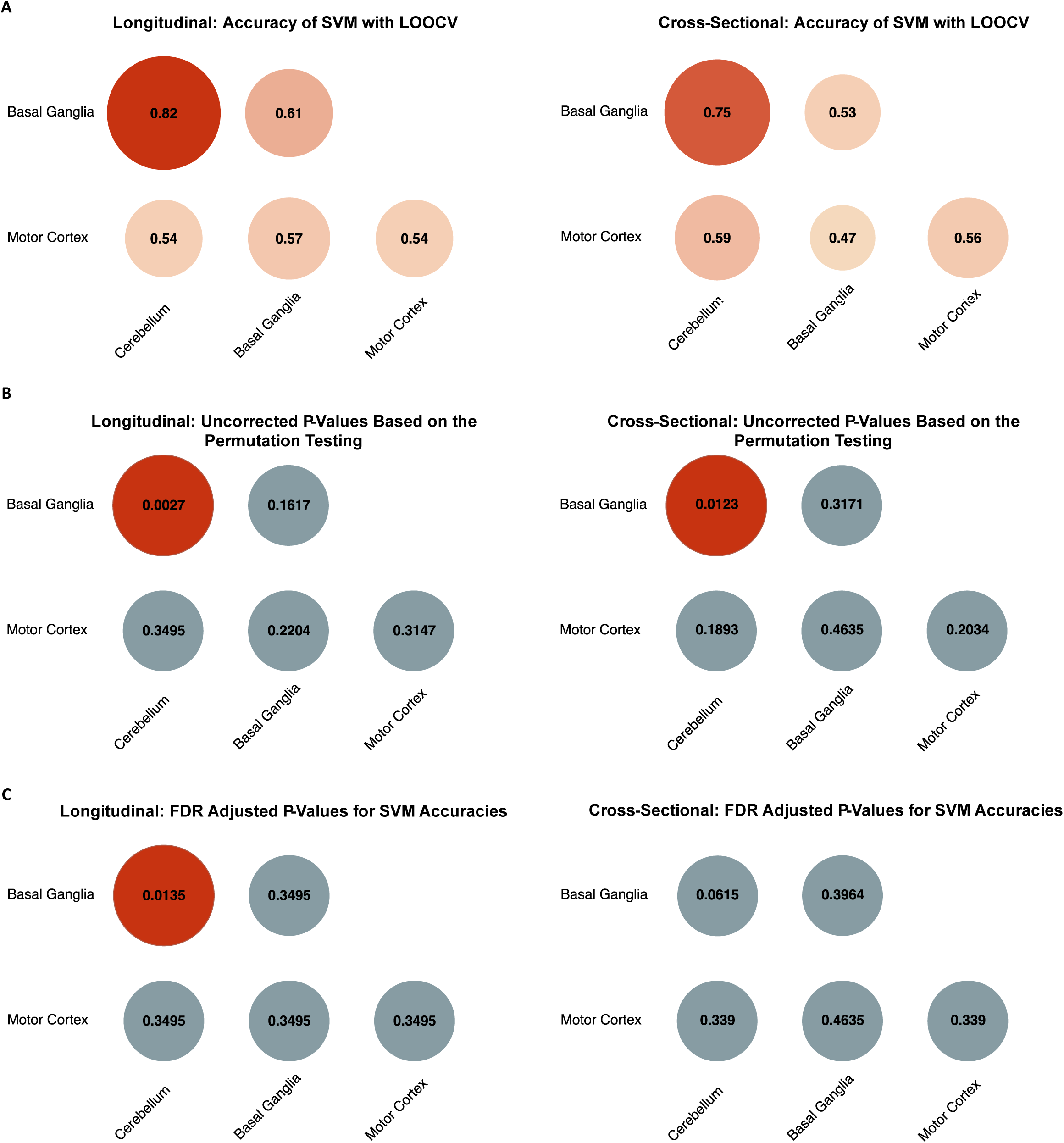
Classification accuracies and statistical significance of support vector machine classifiers. (A) Classification accuracies of linear support vector machine implemented with patterns of connectivity within and across cortical motor, basal ganglia, and cerebellum networks; size of the circles is proportional to the accuracy. (**B)** Uncorrected p-values of each network-level classifier. (**C)** FDR-adjusted p-values of each network-level classifier (significance cut-off = 0.025).

**Figure 5.**
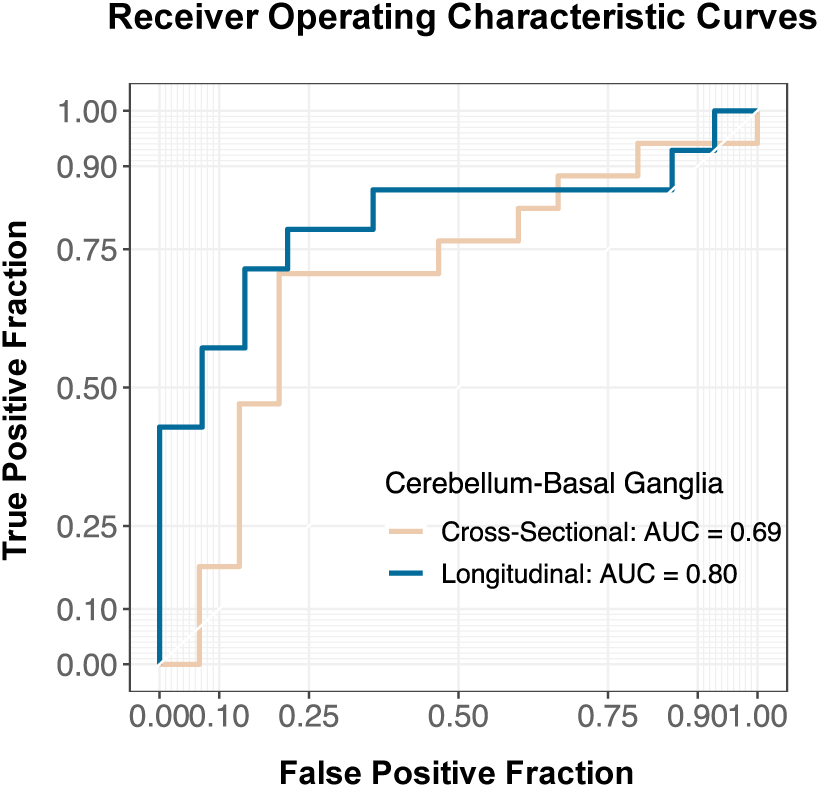
Receiver operating characteristic curve and area under the curve. Receiver operating characteristic curve was constructed for support vector machine using the cerebellum-basal ganglia feature set in both within-subject (blue) and between-group (cream) analyses.

To ensure that the classification of medication status based on BG-cerebellum features was not driven by disease severity, we conducted additional analyses to examine the correlation between clinical scores and SVM classification scores (distance to the decision boundary in the feature space) in each dataset. Our results indicate that there was no statistically significant correlation between these measures. In the longitudinal group, we observed correlation coefficients of -0.14 (p = 0.45), -0.16 (p = 0.38), -0.19 (p = 0.30), 0.09 (p = 0.64), and 0.01 (p = 0.94) for UPDRS-III scores (ON score in medicated state), tremor, bradykinesia/rigidity, gait subscores, and Montreal Cognitive Assessment (MoCA; Nasreddine et al., 2005) scores, respectively. In the cross-sectional patients, we found correlation coefficients of -0.25 (p = 0.21) for UPDRS-III, -0.21 (p = 0.30) for tremor, -0.21 for bradykinesia/rigidity, 0.07 (p = 0.74) for gait, and -0.40 (p = 0.08) for MoCA. These results suggest that our classification approach based on BG-cerebellum features was not influenced by disease severity.

## Discussion

This study aimed to examine the long-term effect of levodopa medication in PD on neural synchrony within the motor CSTC circuitry through cross-sectional and longitudinal datasets. Our result suggests a significant modulatory effect on BG-cerebellum connectivity. Importantly, this effect was evident on network-level functional connectivity and not a single connection. A standard seed-based inter-regional (ROI-to-ROI) functional connectivity analysis, yields no statistically significant effect of the long-term medication on either of the datasets. Yet, in the longitudinal cohort, we observed trends of attenuated FC between cerebellum-motor cortex and within-BG connectivity and elevated GP-motor cortex coherence following levodopa medication. At the network level, the compressed feature set of BG-cerebellum connectivity was significantly different following medication in the longitudinal dataset and exhibited a similar trend in the cross-sectional dataset.

### 4.1 Medication effect on symptom severity

The magnitude of motor response to levodopa is driven by both short-duration response (SDR) and long-duration response (LDR). While SDR is closely related to levodopa plasmatic pharmacokinetics, LDR is a sustained motor improvement induced by chronic levodopa therapy that slowly develops after treatment initiation and lasts days if discontinued (Nutt and Holford, 1996; Wider et al., 2006). Compared to the drug-naïve state, we found less severe overall motor symptoms and bradykinesia in both datasets and additionally milder tremor in the longitudinal analysis at the ON-levodopa state. This result is largely masked by the SDR, where the immediate symptom alleviation roughly parallels the plasma levodopa level. In the longitudinal group, we also observed that clinical measures at drug-naïve and OFF-levodopa states were not statistically distinct, with a median interval of 34.5 (*IQR* = 11 months) months between the two timepoints and a median treatment duration of 22 (*IQR* = 10 months) months. This aligns with the notion that the symptomatic effects of levodopa may delay the natural progression of motor disability through complex mechanisms of LDR (Cilia et al., 2020). Cilia et al. (2020) showed a 31% lower annual decline of UPDRS-III scores in the OFF-levodopa state over a 24-month treatment duration relative to the natural progression (consecutive drug-naïve) of motor symptoms.

### 4.2 Network-level changes in functional connectivity

The SVM-based classification analysis pointed to the relevance of cerebellum-BG network connectivity in discriminating PD medication status (de novo versus levodopa-medicated). The interaction between the cerebellum and BG has traditionally been thought to occur at the cortical level, as both areas form loops with the cortex (for a review see Haar and Donchin, 2020). However, recent anatomical tracing studies have demonstrated bidirectional pathways facilitating a direct cerebellum-BG interplay, which could lead to the formation of an integrated functional network (Bostan and Strick, 2010; Hoshi et al., 2005; Pelzer et al., 2013). Accordingly, resting state striatum-cerebellum functional coupling has been indicated in healthy adults, where the connections are segregated based on functional topography and decline in advanced age, where dopamine has a normative decline (Bernard et al., 2012; Hausman et al., 2020). Further, levodopa was shown to enhance the connectivity in motor pathways joining putamen with the cerebellum and brainstem in healthy individuals (Kelly et al., 2009). These findings led to the speculation that dopamine may be particularly important in modulating the functional interaction between the cerebellum and BG.

In PD, heterogenous results were reported. A recent study by Bagarinao et al. (2022) demonstrated significantly impaired connectivity between cerebellar connector hubs and BG in PD, with altered connectivity correlating with clinical manifestations. Hacker et al. (2012) found markedly lower connectivity between the cerebellum with caudate and putamen in advanced, medicated patients relative to matched controls. In contrast, Helmich et al. (2010) found no difference in striatum-cerebellum connectivity between de novo or off-medication patients and healthy controls. Gao et al. (2017) reported decreased cerebellar connectivity with putamen and GP in off-levodopa patients and subsequent normalisation following levodopa administration. Simioni et al. (2016) described a contrasting pattern of increased coherence between putamen and motor cerebellum in the off- medication mild-moderate patients and the normalising effect of levodopa. The inconsistent results could be due to the differential role of cerebellum-BG interaction at different disease stages and the variations in ROI choice and localisation. It is also possible that the variations originated from a focus on individual connections, and a potentially more coherent network modulation of levodopa might have been missed.

In our study, while pure univariate effects were absent from interregional cerebellum-BG connectivity in differentiating de novo and levodopa conditions, the multivariate combination of individual connections demonstrated robust discriminative power. This indicates a contrast in a distributed pattern of neural synchrony between the medication conditions. Specifically, a network of intra- and cross-hemispheric connectivity between cerebellar and BG nodes acted synergistically to manifest a difference that was not accessible to univariate comparisons. Although we could not rule out the effect of disease progression in the longitudinal dataset, the effectiveness (to a weaker extent) of the feature set in cross-sectional analysis reaffirmed the role of levodopa in altering cerebellum-BG network connectivity patterns in PD. Complementing the previous work investigating connectivity between specific brain regions (Hausman et al., 2020; Kelly et al., 2009), our finding provides new insight into dopamine’s potentially wider, system-level modulation of the reciprocally connected cerebellum-BG network. In PD, increased striatal dopamine following levodopa medication could prevent transmitting aberrant BG signals to the cerebellum that could evoke cerebellar hyperactivity and disrupt cerebello-thalamo-cortical pathways (Bostan and Strick, 2010; Caligiore et al., 2016; Milardi et al., 2019). A refined cerebellum-BG interaction could also facilitate the combination of supervised learning and reinforcement learning (specialised by cerebellum and BG, respectively) and consequently lead to better motor learning and adaptation in PD (Caligiore et al., 2016).

Finally, the levodopa-induced modification of cerebellar-BG connectivity found here concurs with previous works that found a similar effect on this pathway following effective deep brain stimulation (DBS) therapy in PD. deep brain stimulation (DBS). Kahan et al. (2019) conducted dynamic causal modelling in DBS patients and showed that only during a motor task (but not during rest) did the DBS modulate the cerebellum-BG connectivity. A more recent investigation of STN DBS in PD highlighted the importance of cerebellum and cerebellum-BG interaction in PD symptom alleviation (Chu et al., 2023). This study found that DBS modulate the functional architecture of large-scale brain networks, including the restoration of lowered dynamic FC between the cerebellum and BG and motor thalamus (Chu et al., 2023).

### 4.3 ROI-level changes in functional connectivity

The longitudinal trends of ROI-level functional reorganisation were likely confounded by the pathological progression, which was only partially controlled by age and time elapsed between the two measurements. Three main trends were observed.

First, de novo patients were inclined to exhibit higher cerebellum-motor cortex connectivity. Cerebellar hyperactivations in PD were consistently reported during the execution of various upper limb movements and motor learning (Rascol et al., 1997; Wu et al., 2010). In a PET study, early PD patients showed additional activations of the bilateral cerebellum while achieving equal performance in motor sequence learning as the control subjects (Mentis et al., 2003). In accordance with our results, Wu et al. (2009) showed that levodopa normalises the heightened rs-FC in the cerebellum and primary motor cortex in PD. The physiological significance of strengthened connectivity and hyperactivity of the cerebellum might be interpreted as compensatory mechanisms in response to impaired subcortical-cortical loops. It was suggested that by restoring the motor circuits with levodopa, patients might become less reliant on the compensatory effects (Wu et al., 2009).

Second, we observed a trend of attenuated rs-FC among the thalamus, putamen, and GP in longitudinal patients at the second time point. Contrary to our observation, previous studies have shown an acute levodopa effect on improving the deficient rs-FC among BG and thalamus nodes (Gao et al., 2017; Szewczyk-Krolikowski et al., 2014). In a longitudinal investigation, Li et al. (2020) reported a time-related decline in BG connectivity focused on putamen in PD, which was associated with changes in nigrostriatal dopaminergic integrity. Hence, our results could represent the net effect of medication and progressing dopaminergic dysfunction that cannot be effectively disentangled.

Third, longitudinal patients tend to exhibit elevated connectivity between GP and motor areas, including supplementary motor area, premotor, and primary motor cortices. A leading hypothesis of PD pathophysiology is the imbalance between direct and indirect BG pathways due to DA deficiency. Suppression of GP interna (GPi; BG output nucleus) and stimulation of GP externa (GPe) firings in PD were shown to alleviate the motor symptoms, which was presumably due to selective inhibition of the indirect pathway (Assaf and Schiller, 2019; Mastro et al., 2017). Studies in pallidotomy and GPi deep brain stimulation (DBS) patients have shown that metabolic activities in frontal motor areas were reinstated after the procedures, indicating the excessive pallidal inhibition of the thalamocortical system in PD (Fukuda et al., 2001; Grafton et al., 1995; Samuel et al., 1997). Considering the above, our findings might reflect a modulatory effect of levodopa on BG pathways, resulting in enhanced GPe-motor cortex connectivity and attenuated GPi-cortex anti-coherence. Nevertheless, a clear inference cannot be drawn as we did not differentiate GPi and GPe despite their distinctive roles in the BG circuitry.

### 4.4 Limitations

This study has several limitations. First, the causal influence of levodopa on clinical and connectivity characteristics cannot be established with an observational design. Second, no rs- FC contrasts survived after implementing the FDR approach to prevent the occurrence of type I errors. This could be due to the inadequate sample size and, therefore, a lower statistical power. In cross-sectional analysis, the disease duration was relatively more heterogeneous among de novo patients; this within-group heterogeneity might have diluted the evidence for group-level inferences. Third, given that levodopa patients were ON medication during fMRI scans, our results reflected a mixture of short- and long-duration medication effects on functional reorganisation in the brain that can not be disentangled. Fourth, although global signal (GS) reduction during the rs-fMRI pre-processing efficiently excludes the non-neural confounds, it could also remove meaningful neuronal fluctuations and introduce artefacts, which might vary for individual datasets (Chen et al., 2012; Murphy and Fox, 2017). Finally, the SVMs models were not validated through out-of-sample (unseen data) predictions; therefore, the full generalisability of the models cannot be evaluated.

### 4.5 Conclusions

To conclude, we demonstrated the modulatory effect of levodopa on resting-state functional connectivity between the cerebellum and BG networks. This effect was absent in the univariate comparisons of individual inter-ROI connectivity, indicating that levodopa modulated collective patterns of BG-cerebellum neural synchrony. Following the recent evidence suggesting the bidirectional linkage between the cerebellum and BG networks, our results provide further insight into the relevance of the inter-network functional connectivity in Parkinson’s, as well as in the brain functional reorganisation processes

## Materials and methods

### 5.1 Participants

The clinical and rs-fMRI data were extracted from the Parkinson’s Progression Marker Initiative (PPMI; see https://www.ppmi-info.org/), a muti-centre observational study of clinical and neuroimaging progression markers of PD conducted since 2010. The PPMI study recruited PD subjects who (1) had at least two of the cardinal motor symptoms (resting tremor, bradykinesia, and rigidity); (2) had a Hoehn and Yahr progression score of 1 or 2; (3) were not expected to initiate PD medication within 6 months from baseline; (4) were diagnosed within 2 years before the entry and aged at least 30 years at the time of diagnoses. Patients were excluded if they (1) had taken levodopa, dopamine agonists, monoamine oxidase-B inhibitors, or amantadine within 60 days to baseline; and (2) have taken levodopa or dopamine agonists for longer than 60 days prior to baseline.

At the time we downloaded the data, a total of 113 patients had had at least one rs-fMRI scan. Here, we aimed to derive a cross-sectional cohort comprising two independent arms of de novo and levodopa-medicated patients. Pertinent thresholds were set for disease and treatment durations (at the time of the scan) based on overall distributions to ensure the former was comparable between the groups and the latter was distributed with a low deviation value in the levodopa group (de novo group: N = 15; levodopa group: N = 17). Moreover, we extracted a longitudinal group where the patients had had at least two rs-fMRI scans, first during the drug-naïve phase and second after transitioning into the levodopa-medicated state (co-administered with a dopa-decarboxylase inhibitor; not on other PD medications), with a moderately concentrated distribution of treatment durations at the second visit across the samples (N = 14).

### 5.2 Clinical measurements

All subjects underwent motor and neuropsychological examinations at the time of selected visits. The presence of motor signs was evaluated using Movement Disorder Society-Unified Parkinson’s Disease Rating Scale (MDS-UPDRS; Goetz et al., 2008). We subdivided the UPDRS scale into a tremor score (items 2.10 and 3.15-3.18), a bradykinesia and rigidity score (items 3.3-3.8), as well as a gait disturbance score (items 2.12, 2.13, 3.10, 3.11, and 3.12) based on previous studies (Aleksovski et al., 2018; Ng et al., 2017). For the non-motor dimension, we included the Geriatric Depression Scale (GDS; Sheikh and Yesavage, 1986) and Montreal Cognitive Assessment (MoCA; Nasreddine et al., 2005).

### 5.3 Imaging acquisition, pre-processing, and region-of-interest selection

#### 5.3.1 MRI acquisition

Whole-brain T1-weighted anatomical MRI and rs-fMRI scans using Siemens Trio Tim 3 Tesla magnets (Siemens Medical Solutions, Erlangen, Germany) were acquired from the PPMI database. Imaging parameters were identical across the clinical sites. Briefly, T1-weighted high-resolution anatomical image (voxel size: 1×1×1 mm^3^) was obtained for each patient with repetition time (TR) = 2,300 ms, echo time (TE) = 2.98 ms, flip angle (FA) = 9 °. rs-fMRI echo-planer scans were conducted in 8.5 minutes with 210 time points, TR = 2,400 ms, TE = 25 ms, FA = 80 °, voxel size = 3.25×3.25×3.25 mm^3^.

#### 5.3.2 Anatomical and functional pre-processing

For the cross-sectional dataset, anatomical reconstruction (cortical) and segmentation (subcortical) were performed using FreeSurfer (version 7.1.1; https://surfer.nmr.mgh.harvard.edu/). The processes included motion correction, removal of non-brain tissues, intensity normalisation, and classification of voxels into white matter (WM) and grey matter (GM) based on intensity and neighbour constraints. In the surface-based stream, the WM and pial surfaces were constructed and refined to follow the intensity gradients between WM and GM and between GM and cerebrospinal fluid, respectively (Fischl and Dale, 2000). Individual surfaces were then aligned to the spherical Destrieux atlas through a non-linear registration algorithm, which fitted cortical folding patterns to an average cortical geometry with each hemispheric cortex parcellated into 74 regions (Destrieux et al., 2010; Fischl et al., 2001). In the volume-based stream, the subcortical regions were segmented and labelled in the native space (Fischl et al., 2004, 2002). For the longitudinal dataset, FreeSurfer longitudinal pipeline was used to construct an unbiased subject-specific template through an inverse consistent registration between the two scans (Reuter and Fischl, 2011). The common information on within-subject templates then initialised the image processing steps (see above) for scans at each visit (Reuter et al., 2012).

Rs-fMRI pre-processing was conducted using FreeSurfer FS-FAST (https://surfer.nmr.mgh.harvard.edu/fswiki/FsFastTutorialV6.0/FsFastPreProc/). The procedure entailed normalizing intensity and generating a mean global waveform (subsequently used as a nuisance regressor), motion correction, and slice timing correction. Pre-processed anatomical volumes were resampled into native rs-fMRI space using mri_vol2vol and bbregister to extract the corresponding time courses of segmented regions.

#### 5.3.3 ROI selection

We defined subcortical and cortical ROIs comprising the motor cortico-striatal-thalamic-cortical (CSTC) circuitry, along with cerebellar ROIs (Table 1). The motor ROI cluster included bilateral precentral gyri (PreCG), inferior precentral sulci (InfPreCS), superior precentral sulci (SupPreCS), caudal superior frontal gyri (SFGcau; posterior 1/3 of superior frontal gyrus), and caudal middle frontal gyri (MFGcau; posterior 1/3 of middle frontal gyrus) based on the Destrieux atlas parcellation (Destrieux et al., 2010). PreCG, InfPreCS, and SupPreCS were localised in the main body, anteroinferior, and anterosuperior subsections of the primary motor cortex, respectively. SFGcau corresponded to the anatomical location of the supplementary motor area, and MFGcau was functionally mapped to premotor cortices. At the subcortical level, we focused on the bilateral thalamus, caudate nucleus, putamen, globus pallidus, and cerebellar cortex.

**Table 1.** Selected ROIs.

|  |
| --- |
| **Subcortical ROIs (voxels)** |
| Cerebellum cortex (CB) |
| Thalamus (Tha) |
| Caudate nucleus (Cdn) |
| Putamen (Pu) |
| Globus pallidum (GP) |
| **Motor cortical ROIs (vertices)** |
| Precentral gyrus (PreCG) |
| Inferior precentral sulcus (InfPreCS) |
| Superior precentral sulcus (SupPreCS) |
| Caudal superior frontal gyrus (SFGcau) |
| Caudal middle frontal gyrus (MFGcau) |

Time courses of subcortical ROIs are voxel-based, whereas the fMRI data of motor cortical ROIs are vertex-based.

### 5.4 Statistical analysis

#### 5.4.1 First-level analysis of rs-fMRI data

All analyses were conducted using MATLAB R2021a (https://uk.mathworks.com/products/matlab.html). Voxel- (volume space) and vertex-level (surface space) time-series data for each ROI, cerebrospinal fluid (CSF), and WM were extracted, along with the whole-brain average time course. The first four images of the fMRI time-series were discarded. To exclude spurious signal confounds, we implemented a stepwise noise attenuation procedure. First, a high-pass filter with a threshold of 0.01 Hz was used, and the six head motion parameters estimated by FreeSurfer were regressed out of all raw fMRI data through linear regression. The voxel-level time courses were averaged for CSF and WM, then regressed out from the global time course and all voxels and vertices that constitute each ROI. Next, nuisance regressions of the global wave to voxel/vertex-level time series were conducted, and the cleaned residuals from the fits were averaged to create ROI time traces. Lastly, before mean-centring and standardisation, we further denoised the ROI time series with a low-pass filter retaining frequencies below 0.20 Hz to remove the potential high-frequency fluctuations injected in partialling-out processes. The pairwise Pearson correlation coefficients between the filtered time-series of each ROI were computed and stored in a 20*20 symmetric FC matrix.

#### 5.4.2 Second level within- and between-group comparison

Nonparametric permutation testing was used to extract significant differences in the connectivity maps of patients in de novo and levodopa-medicated states in both cross-sectional (between-group) and longitudinal (within-group) datasets. The permutation inferences assume that data can be arbitrarily exchanged without affecting the joint probability distribution (Winkler et al., 2014). Nevertheless, given the presence of nuisance covariates, such as disease duration, age, and gender, the data cannot be considered exchangeable even under the null hypothesis. In this respect, the Freedman-Lane procedure was followed to compute the estimates of null distribution to ascribe p-values for each rs-FC (Freedman and Lane, 1983; Winkler et al., 2014; Zalesky et al., 2010). First, we fitted a full general linear model (GLM):

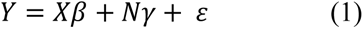

For the cross-sectional dataset, *Y* included the pairwise Pearson’s *r*, regressor *X* contained group identity, and *N* incorporated disease duration, age, and gender. For the longitudinal dataset, *Y* included the within-subject contrast of each connectivity (levodopa – de novo), *X* was a vector of ones (the overall mean effect), and *N* contained the difference in disease duration between two timepoints, baseline age, and gender. Parameters *β* and *γ* were, respectively, for factors of interest and nuisance variables, and *ε* represented the residual. We then created a reduced GLM by regressing *Y* only with nuisance variables and obtained estimated regression parameters *γ̂*_*N*_ and nuisance-only residuals *ε̂*_*N*_ [variance of the null model (Equation 2)]. Here, the observed F-statistic *F*_0_ (based on sums of squares in unpermuted GLMs) in the cross-sectional dataset and the mean connectivity difference *Y̅*_0_ in the longitudinal dataset were derived as test-statistics *T*_0_.

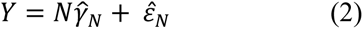

Next, we shuffled the residual vector through multiplying by a permutation matrix *P*_*j*_. The nuisance signal estimated in the earlier step, *Nγ̂*_*N*_, was then added back to the permuted residuals, *P*_*j*_*ε̂*_*N*_, to produce the permuted estimations of connectivity strengths (cross-sectional) and contrasts (longitudinal) *Y*_*j*_:

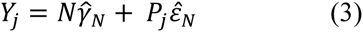

Finally, the permuted estimates were regressed against the full model (Equation 4), including both primary and nuisance factors, and the test-statistics *T*_*j*_ were computed.

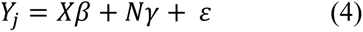

The permutation was repeated 10,000 times to generate the null distribution of test statistics. Each FC was deemed significantly different (uncorrected) between the medication states if the absolute observed *T*_0_ was greater than 95% of the absolute values of distributed *T*_*j*_. Finally, the false discovery rate (FDR)-based multiple comparison procedure was adopted to adjust the derived p-values (Benjamini and Hochberg, 1995; Yekutieli and Benjamini, 1999).

#### 5.4.3 Network level testing of medication effect on functional connectivity

To evaluate the discriminating power of neural synchrony at the network level, we consider three subnetworks of the motor CSTC circuitry: cortical motor areas, sub-cortical motor areas (BG and thalamus), and cerebellum. For all three subnetworks, we integrated the hemispheres. We then constructed support vector machines (SVM) implemented on within- and cross-network features to classify patients with respect to medication status. Both the feature and classifier constructions were conducted for the two datasets separately. We first took all pairwise combinations of the networks (discarding the within-cerebellum coupling due to only one ROI-level feature) as masks, and subject connectivity values in each mask were collapsed into a feature vector. To this end, five feature vectors (one for each of the within and between networks connections) with varying lengths (according to the number of ROIs in each network) were retrieved, with the numbers of FC values in most vectors outnumbering the number of observations.

##### 5.4.3.1 Dimensionality reduction

To attenuate the risk of overfitting and ensure the predictive algorithms based on different network-level feature sets are comparable, we used principal component analysis (PCA) to linearly transform correlated FC values into a reduced number of orthogonal variables, i.e., principal components (PC), to derive new vector sets with the same dimensionality (Jolliffe, 2002). In specific, eigen decomposition of the covariance matrices from each standardised feature set was performed to compute eigenvalues and eigenvectors. The eigenvalues were then ranked in a descending order effectively representing decreasing variance in the data carried by the corresponding PCs, whose directions were represented by the associated eigenvectors (Jolliffe, 2002; Mwangi et al., 2014). For each group of FC values, the number of PCs was determined using a 99.5% threshold on the cumulative percentage of total variance.

##### 5.4.3.2 Support vector machine and cross-validation

Binary classifications of de novo and levodopa-medicated patients were performed using SVMs with the linear kernel through the *fitcsvm* route in MATLAB. Detailed documentation of SVM and the optimisation problem can be found in Cortes et al. (1995). Briefly, a linear SVM projects the training data points into a high-dimensional feature space and seeks an optimal hyperplane with the maximal margin separating the two classes (Cortes et al., 1995). The optimal hyperplane is defined as:

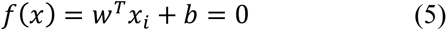

where *w* is the weight vector perpendicular to the hyperplane, *b* is the bias term, *x*_*i*_ is *i*-th input vector in the dataset, and *f*(*x*) is the linear discriminant function whose sign represents the class of training inputs (Cortes et al., 1995). The objective of maximising the geometric margin in the feature space corresponds to the primal optimisation problem (which is then reformulated into the Lagrangian dual function), 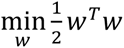, subject to, *y_i_* (*w*^*T*^*x*_i_ + *b*) ≥ 1.

The SVM models were evaluated with leave-one-out cross-validation (LOOCV) for both longitudinal and cross-sectional analyses. In the former, a subsample contained data of the same patient measured at two medication conditions. Permutation testing was applied to assess whether each classifier captured a real class structure in the data. Here, we permuted the class labels and refitted each SVM 10,000 times to estimate the corresponding null distribution of accuracy values. The accuracy was considered significantly higher than chance if it exceeded the 97.5th percentile of the null distribution. For the classifiers with above-chance accuracy, we computed the area under the receiver operating characteristic curve (AUC) that aggregates the classifying efficacy under all possible decision thresholds.

### Data availability

All data used are fully available in the Parkinson’s Progression Markers Initiative (PPMI) database (https://www.ppmi-info.org/)

## Acknowledgements

S.H. is supported by the Edmond and Lily Safra Fellowship. S.A and S.H. are supported by the UK Dementia Research Institute, Care Research & Technology Centre. The data used in the preparation of this article were obtained from the Parkinson’s Progression Markers Initiative (PPMI) database (https://www.ppmi-info.org/). PPMI – a public-private partnership – is funded by The Michael J. Fox Foundation for Parkinson’s Research and funding partners listed here https://www.ppmi-info.org/about-ppmi/who-we-are/studysponsors.

## Funding

No funding was received towards this work.

## Competing interests

The authors report no competing interests.

## Notes

### Competing Interest Statement

The authors have declared no competing interest.

### Summary of Updates

Figures and tables formats

